# Age alters the relationship between post-encoding sleep quality and context memory neural reinstatement

**DOI:** 10.64898/2026.06.17.733023

**Authors:** Masoud Seraji, Soroush Mirjalili, Chuu Nyan, Audrey Duarte, Vince D Calhoun

**Affiliations:** Tri-Institutional Center for Translational Research in Neuroimaging and Data Science (TReNDS), Georgia State University, Georgia Institute of Technology and Emory University, Atlanta, GA, USA; Department of Psychology, University of Texas at Austin, Austin, USA; Department of Psychology, University of Oregon, Eugene, USA

**Author notes:** Co-senior authors contributed equally to this work (Audrey Duarte, Vince D Calhoun).

## Abstract

Sleep supports episodic memory consolidation, yet it remains unclear how naturalistic post-encoding sleep quality relates to the neural reinstatement of episodic representations across adulthood. The present study examined whether sleep discontinuity during the retention interval predicted delayed context memory and encoding–retrieval similarity (ERS) of EEG in younger and older adults. Participants completed an object–scene context memory task with immediate and delayed retrieval, while EEG was recorded during encoding and retrieval. Actigraphy was used to measure sleep across the post-encoding retention period, and principal component analysis identified sleep discontinuity and sleep time components. Behavioral results showed that greater post-encoding sleep discontinuity, but not sleep time, was associated with poorer delayed memory accuracy for mismatching object–context pairs across age. ERS analyses further showed that greater sleep discontinuity was associated with reduced ERS for correctly rejected mismatching pairs across frontal and posterior spatiotemporal clusters. Age moderated sleep–ERS associations: greater sleep discontinuity was generally related to lower ERS in younger adults, whereas some spatiotemporal clusters showed positive associations in older adults, potentially reflecting compensatory or effortful retrieval-related processing in poorer sleepers. Together, these findings suggest that sleep continuity during the post-encoding retention interval is important for preserving high-fidelity episodic representations needed for later context discrimination. More broadly, the results demonstrate that naturalistic sleep fragmentation is linked to both behavioral memory outcomes and neural reinstatement across adults.

## 1. Introduction

Episodic memory depends on the successful encoding, stabilization, and later recovery of event-specific information, including the contextual details that define where and under what circumstances an event occurred. A large literature indicates that sleep contributes to these processes, perhaps most notably the consolidation of newly encoded memories (Diekelmann & Born, 2010; Rasch & Born, 2013). Many studies using sleep deprivation methods suggest that rather than passively protecting memories from interference, post-encoding sleep may facilitate the reorganization and strengthening of hippocampus-dependent traces in ways that support later retrieval (Diekelmann & Born, 2010; Rasch & Born, 2013). These consolidation mechanisms may be especially important for remembering not only an item but its event-specifying associations (for review see, Diekelmann et al., 2009), which depend upon hippocampal relational binding mechanisms (Diana et al., 2007).

Although they allow for the necessary contribution of sleep to be investigated, sleep deprivation manipulations do not enable the measurement of naturalistic sleep in one’s home environment. These studies are typically limited to a single night, precluding the ability to measure night-to-night variability in sleep. A common approach to assessing naturalistic sleep over multiple nights is actigraphy, which uses wearable devices to monitor body movements that indicate sleep-wake behavior. Numerous studies using this approach have shown that even healthy aging, in the absence of diagnosed sleep or cognitive dysfunction, is accompanied by reduced sleep quantity and quality, as indicated by more fragmented sleep, and more time spent awake after sleep onset (Carlson et al., 2023; Mander et al., 2017; Scullin & Bliwise, 2015), as well as greater night-to-night variability in these measures (Hokett et al., 2022; Shoji et al., 2015). Concomitant with declines in sleep quality with increasing age are those in episodic memory, particularly for the associations among elements of an event, including the contextual associations in which items are encoded, rather than memory for individual items (Bender et al., 2010; Craik & Rose, 2012; Naveh-Benjamin et al., 2004). Interestingly, as noted above, memory for these kinds of associations may be particularly dependent upon sleep in younger and older adults, as shown primarily in sleep deprivation studies (for review see, Diekelmann et al., 2009). Emerging findings using actigraphy to monitor naturalistic sleep quality have shown that sleep quality, including average sleep continuity and night-to-night stability therein, may be more strongly tied to episodic memory than sleep duration, especially in adulthood and aging (Hokett et al., 2021; Wilckens et al., 2014). In our systematic review and meta-analysis, we showed that individual differences in naturalistic sleep quality are reliably associated with episodic memory performance across younger and older adults (Hokett et al., 2021). Other studies have shown that these associations may be even stronger for older than younger individuals (Hokett et al., 2022; Wilckens et al., 2014). Collectively, these results suggest that reduced sleep quality may be a contributing factor to age-related episodic memory impairment, although the precise behavioral and neural consequences of these sleep quality reductions remain incompletely understood (Mander et al., 2017; Scullin & Bliwise, 2015).

Neurocomputational models suggest that successful episodic memory retrieval is dependent upon neural reinstatement during retrieval of neural patterns and associated cognitive processes engaged when an event was first encoded (Norman & O’Reilly, 2003). This idea has been supported by fMRI and EEG studies using encoding–retrieval similarity (ERS) approaches, which quantify the extent to which retrieval-related neural activity correlates with encoding-related patterns (Waldhauser et al., 2016; Wing et al., 2015; Xiao et al., 2017). More broadly, this approach is consistent with a growing shift in EEG analysis from isolated component amplitudes toward multivariate, time-resolved approaches that relate distributed neural patterns to behavior (Eichele et al., 2009). These studies have shown greater ERS for successfully remembered events than for forgotten events, above and beyond similarity attributable to shared category or modality features (e.g. Ritchey et al. 2013). Emerging studies using both fMRI and EEG show that aging is associated with reduced ERS, which is associated with age-related reductions in episodic memory performance (Chamberlain et al., 2022; Lee et al., 2022). These findings support the idea that a reduction in neural reinstatement of encoding-related activity patterns may be a mechanism underlying age-related reductions in event-specifying memory details.

We have recently shown that lower naturalistic sleep quality, specifically fragmented sleep, is associated with altered patterns of oscillatory EEG power ERS across the adult lifespan (Hokett et al., 2022). Specifically, participants were asked to encode word pairs that were presented either intact or rearranged in the same day retrieval test. Better average sleep quality measured over the week preceding performance of the memory test was associated with a superior ability to correctly recognize intact and correctly reject rearranged pairs, especially in older age, and greater ERS for rearranged pairs, across age. The ability to correctly reject rearranged pairings is thought to be dependent upon the recollection of prior, correctly paired, associations in order to reject rearranged ones (i.e. “recall-to-reject”) (Rotello, 2000), a cognitive control process negatively impacted by age (Duarte and Dulas 2020; Trelle et al. 2020). These findings are consistent with the idea that recollection-based memories, which are disproportionately impacted by age, may also be particularly sensitive to sleep (for review see, Diekelmann et al., 2009).

Findings from sleep deprivation studies suggest that *post-encoding* sleep quality should be associated with episodic memory and the neural reinstatement of event-specific representations. Yet, the role of naturalistic sleep quality during the delay between encoding and retrieval remains underexplored, especially with respect to event-specific neural reinstatement. Given that we did not impose a delay between encoding and retrieval in our prior study (Hokett et al., 2022), we could not test these predictions. The present study addressed these gaps by measuring sleep quality using actigraphy over a 96-hour period between encoding and retrieval of an object-scene context memory task during which EEG was recorded. With this object-scene context memory task, we are able to determine whether our prior findings for word pairs generalize to other kinds of episodic associations, as would be predicted by theories of sleep-dependent memory consolidation (Stickgold, 2005). Representational similarity analysis was used to measure encoding-retrieval neural reinstatement. With this approach, we could test the hypothesis that worse, more fragmented post-encoding sleep is associated with altered reinstatement of contextual memory representations, particularly under conditions that require discrimination of mismatching context associations, and also test the possibility that these relationships are stronger with increasing age (Hokett et al., 2022; Wilckens et al., 2014). Given that processing linked to recollection and episodic reconstruction as well as ERS have been shown to emerge during retrieval from ∼400-500 ms until response (Hokett et al., 2022; James et al., 2016; Johnson et al., 2015; Lu et al., 2015; Rugg & Curran, 2007), we predicted that ERS in this time period would be sensitive to both sleep and age. This novel study links individual differences in real-world sleep quality during the critical post-encoding period to the temporal dynamics of episodic memory reinstatement and the impact of age on these relationships.

## 2. Materials and Methods

### 2.1. Participants

Ninety right-handed adults were initially recruited for the study. Principal component analysis (PCA) was used to examine sleep-related patterns in the data; participants were included in these analyses if they had usable sleep and imaging measures, 10 participants were excluded because retention-night sleep data were unavailable, including 9 participants with accelerometer malfunction resulting in data loss, and 1 participant who did not wear the device on the retention nights. Thus, 80 participants were included in the sleep PCA analyses. For the EEG analyses, 10 participants were excluded due to participation in an earlier version of the task with different stimulus/trigger coding (n = 4), researcher error in presenting experimental files (n = 2), participant failure to use all of the response keys (n = 2), technical issues resulting in loss of EEG trigger codes (n = 1), and excessive EEG artifacts caused by sweat (n = 1). The final EEG sample consisted of 70 participants, including 38 females, 31 males, and 1 non-binary participant from two age groups: younger adults (18–30 years) and older adults (56–79 years). All participants were native English speakers with normal or corrected-to-normal vision and no reported hearing impairments. Handedness was verified using the Edinburgh Handedness Inventory. Prior to participation, individuals completed a detailed screening questionnaire to confirm the absence of any psychiatric, neurological, or sleep disorders, as well as any history of cardiovascular or cerebrovascular disease. All participants provided written informed consent. The experimental protocol and recruitment procedures were approved by the Institutional Review Board (IRB) of the University of Texas at Austin. Participants received monetary compensation for their time and effort.

### 2.2. Study Procedure and Timeline

The study followed a seven-day protocol that combined laboratory visits with at-home sleep monitoring (Fig. 1). On Day 0, participants completed an initial laboratory session that included a neuropsychological assessment to screen for mild cognitive impairment and fitting with Actiwatch-2 accelerometers (Philips Respironics) to monitor naturalistic sleep–wake patterns. They were asked to wear accelerometers continuously at home on Days 1–2, prior to the memory task. On Day 3, participants returned to the lab for the encoding phase followed immediately by an immediate retrieval test. Actigraphy recording continued during Days 4–6, allowing characterization of post-encoding sleep (i.e. retention sleep). Finally, on Day 7, participants completed a delayed retrieval session and returned the accelerometers.

**Fig. 1.**
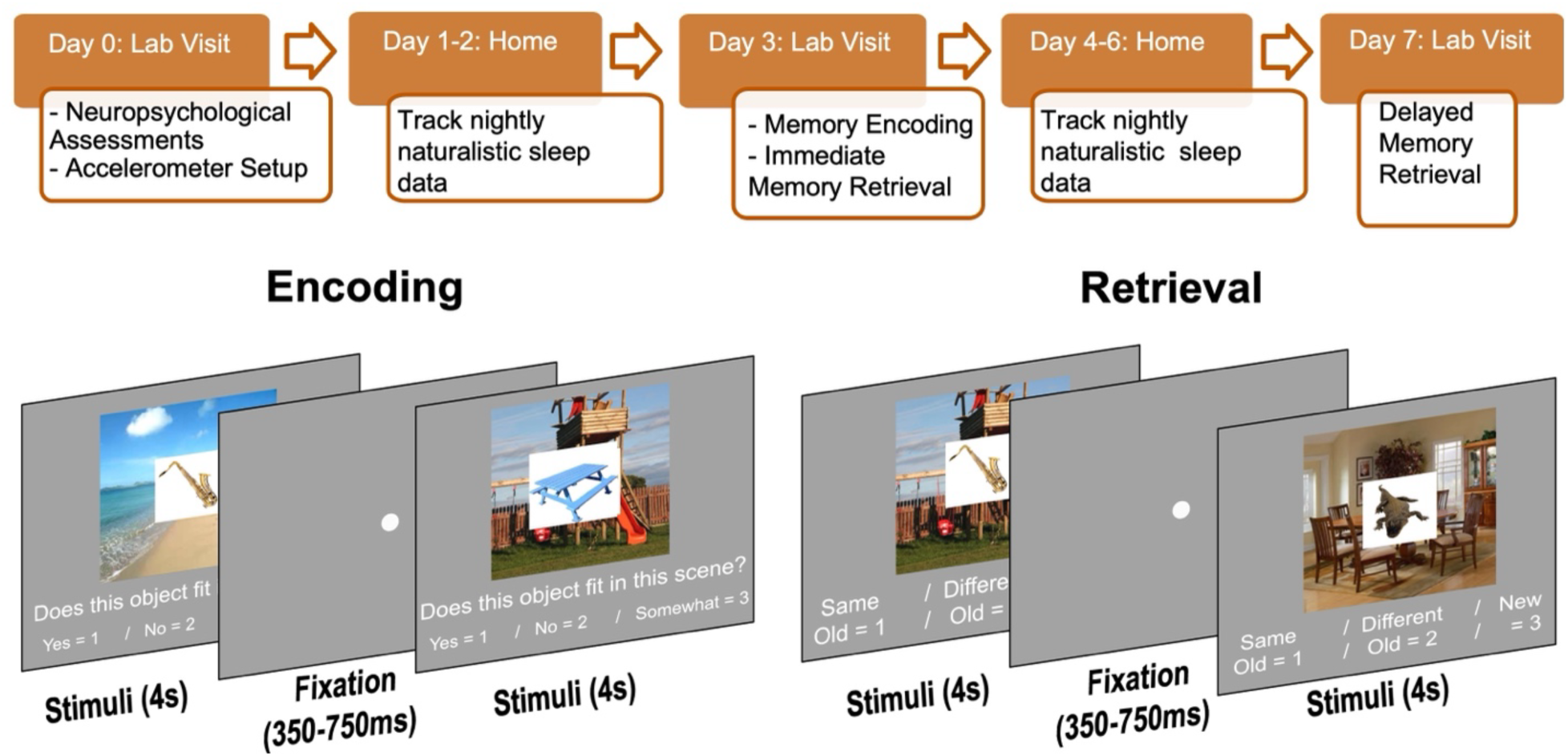
Study timeline and memory task structure. Overview of the seven-day experimental protocol. On Day 0, participants completed neuropsychological assessments and were fitted with an Actiwatch-2 accelerometer. Days 1–2 involved at-home tracking of naturalistic sleep. On Day 3, participants returned to the laboratory for the memory session, which included the encoding task followed by an immediate retrieval test. Participants continued sleep monitoring at home during Days 4–6. On Day 7, they completed a delayed retrieval session. The bottom panels illustrate the trial structure for the encoding (left) and retrieval (right) tasks. During encoding, each trial consisted of an object–scene image for which participants judged whether the object fit (suited) within the scene. During retrieval, participants were asked to decide whether the object presented was old and previously presented with the same scene (“Same, Old”) old but not previously presented with the scene shown (“Different, Old”) or “New.”

During encoding on Day 3, participants studied 288 unique full-color objects presented on one of 12 scenes. Object images were drawn from two sources: (1) the Hemera Photo-Objects image library and (2) additional high-quality photographs obtained from Google Images, selected for clarity and recognizability. Scenes were sampled from 12 distinct categories spanning outdoor environments (e.g., beach, park, forest) and indoor settings (e.g., kitchen, office, living room). Each trial began with a jittered fixation cross (350–750 ms), followed by a 4-s presentation of the object embedded within the scene. Participants judged whether the object logically fit within the scene using three response options: “Yes” (1), “No” (2), or “Somewhat” (3).

Participants completed two retrieval phases—an immediate retrieval on Day 3 and a delayed retrieval on Day 7. Each retrieval session consisted of 432 trials, comprising the 288 previously studied objects (“old” trials) intermixed with 144 new object images drawn from the same two sources used at encoding (Hemera Photo-Objects and curated Google Images). New objects were paired with background scenes sampled from the same 12 scene categories used during encoding, ensuring the contextual backgrounds remained familiar while the objects were novel. On each retrieval trial, participants viewed a fixation followed by a 4-s object-scene image and categorized the stimulus into one of three conditions: “Same Old” (object and scene identical to encoding), “Different Old” (previously seen object presented with a different scene), or “New” (novel object paired with a familiar scene). Immediate and delayed retrieval differed primarily in the timing of the test. Immediate retrieval was completed shortly after encoding, whereas delayed retrieval was completed on Day 7. The old-item conditions were matched across the two retrieval sessions: intact trials contained the same object–scene pairs, and recombined trials contained the same object–scene rearrangements as in immediate retrieval. In contrast, novel-object trials were not repeated across retrieval sessions. A separate set of new objects was used during delayed retrieval so that these items remained novel at the time of the delayed test.

### 2.3. Actigraphy-derived sleep metrics and dimensionality reduction

Actigraphy recordings were used to quantify nightly sleep behavior across the seven-day monitoring period. Our primary interest was in the 4 nights of post-encoding sleep. From these nights, we extracted five variables: total sleep time (TST), sleep efficiency, sleep fragmentation, wake after sleep onset (WASO), and number of wake bouts. TST represented the total minutes scored as sleep. Sleep efficiency indexed the percentage of time in bed spent asleep. Sleep fragmentation quantified the extent of nighttime sleep disruption, indexing repeated awakenings and interruptions that reduced sleep continuity across the night. WASO captured the cumulative minutes spent awake after initial sleep onset, and wake bouts reflected the number of discrete awakenings throughout the night. For each participant, we computed the across night means for these variables. To reduce dimensionality and account for shared variance among sleep measures, we conducted a PCA on mean sleep metrics derived from actigraphy in participants with complete data (n = 80). The PCA was performed on standardized (z-scored) variables using a Varimax rotation to enhance interpretability by maximizing the separation of variable loadings across components. Component retention was determined using the Kaiser criterion (eigenvalues > 1). Component scores were standardized and used as continuous predictors in the hierarchical regression models relating sleep behavior to encoding–retrieval similarity and memory performance.

### 2.4. EEG acquisition and processing

Continuous scalp-recorded EEG data were collected during encoding and both retrieval phases using 31 active Ag/AgCl electrodes mounted on a BrainVision ActiCAP system (Brain Products GmbH, Germany), following the extended 10–20 international placement system (Nuwer et al., 1998). Electrodes were positioned at the following sites: Fp1, Fz, F3, F7, FT9, FC5, FC1, C3, T7, TP9, CP5, CP1, Pz, P3, P7, O1, Oz, O2, P4, P8, TP10, CP6, CP2, C4, T8, FT10, FC6, FC2, F4, F8, and Fp2. Cz was used for online referencing. Horizontal and vertical electrooculogram (HEOG and VEOG) signals were recorded using four external electrodes placed at the outer canthi and above/below the right eye to monitor eye movements and blinks. EEG signals were continuously sampled at 500 Hz with 24-bit resolution, and no online filtering was applied during acquisition. Electrode impedances were maintained below 10 kΩ throughout the experiment. Offline preprocessing was performed in MATLAB (The MathWorks Inc., Natick, MA) using the EEGLAB (Delorme & Makeig, 2004) and FieldTrip(Oostenveld et al., 2011) toolboxes. Data were down-sampled to 250 Hz and re-referenced to the average of the left and right mastoid electrodes (TP9 and TP10). A band-pass filter of 0.05–80 Hz was applied, and 60 Hz line noise was removed. Continuous data were then segmented into stimulus-locked epochs spanning –1,000 to 3,000 ms relative to image onset for all encoding and retrieval trials (covering all 288 encoding and 432 retrieval events). Each epoch was baseline-corrected using the –400 to –200 ms pre-stimulus interval, ensuring a common reference period for both encoding and retrieval analyses.

Artifact rejection was carried out in multiple stages to maximize signal quality. First, channels and epochs exhibiting excessive drift, clipping, or flatlining were flagged using automatic criteria, including voltage fluctuations exceeding ±150 μV within a trial. These segments were removed or, in the case of isolated bad channels, interpolated from neighboring electrodes. Independent component analysis (ICA) was then applied to the remaining epoched data to isolate and remove components reflecting ocular artifacts, such as blinks, saccades, and other eye-movement–related activity (Delorme & Makeig, 2004). ICA and related source-separation approaches have also been widely used to separate latent electrophysiological processes and artifacts in EEG and multimodal EEG–fMRI data (Eichele et al., 2009). Components were identified based on their scalp topographies, time courses, and power spectra, and those clearly associated with ocular or non-neural artifacts were excluded from back-projection. Following ICA correction, all trials were visually inspected to ensure that residual artifacts were minimized, and overall data integrity was preserved.

Time–frequency decomposition was conducted using complex Morlet wavelet (Tallon-Baudry & Bertrand, 1999) transforms, a method well suited for capturing transient oscillatory activity with high temporal precision. Wavelets were applied at 38 logarithmically spaced frequencies between 3 and 40 Hz, allowing us to characterize low-, mid-, and high-frequency dynamics on a common scale. Prior to decomposition, the data were down-sampled to 50 Hz to reduce computational load while preserving the temporal resolution necessary for analyzing cognitive oscillations. For each trial, wavelet convolution produced power estimates at each electrode and frequency across 120 time points, each corresponding to a 20-ms interval, resulting in a four-dimensional data structure organized as trials × electrodes × frequencies × time points. To examine the temporal evolution of neural oscillations at a broader scale, the time–frequency power matrices were further aggregated into 22 partially overlapping windows, each spanning 300 ms and advancing in 100-ms steps across the post-stimulus period (0–2,400 ms). This sliding-window approach smooths high-frequency fluctuations while capturing gradual changes in oscillatory power across the unfolding cognitive process.

### 2.5. Representational Similarity Analysis of Encoding–Retrieval EEG Patterns

Representational similarity analysis (RSA) was used to quantify the extent to which oscillatory EEG patterns present during encoding were correlated with those during delayed retrieval, potentially reflecting neural reinstatement of encoding processes, and how this reinstatement was related to sleep quality during the post-encoding retention period. Data from immediate retrieval were not included in subsequent analyses. This decision allowed the analyses to focus specifically on delayed memory-related neural reinstatement after the retention interval, when sleep-dependent consolidation processes were expected to have occurred.

To construct feature vectors for RSA, we extracted oscillatory power from 3–40 Hz for every encoding and retrieval trial. For each trial, this yielded a time–frequency matrix of size (38 frequencies × T time points). Oscillatory power was then segmented into 300-ms windows, producing a reduced representation of size (38 frequencies × ∼22-time windows) per trial. Within each time window, power values were log-transformed to reduce skewness. Analyses were then performed separately for four predefined electrode groups based on scalp region: left frontal (FC1, FC5, F3, F7, FT9, FP1), right frontal (FC2, FC6, F4, F8, FT10, FP2), left posterior (CP1, CP5, P3, P7, TP9, O1), and right posterior (CP2, CP6, P4, P8, TP10, O2). This regional grouping reduces the number of statistical comparisons, increases robustness by averaging across nearby electrodes, and preserves meaningful hemispheric and anterior–posterior distinctions relevant to memory-related oscillatory activity. Power was first averaged across frequencies within each window, yielding a (6 electrodes × ∼22-time windows) matrix per trial. We then averaged across electrodes within the cluster. Thus, for every encoding trial and every retrieval trial, each electrode cluster contributed one multidimensional feature vector representing log-transformed oscillatory power across the series of 300-ms windows. These vectors served as the inputs for representational similarity analysis. Fig. 2. illustrates the steps involved in constructing these representations, including the electrode clusters (panel A), time–frequency power matrices for encoding and retrieval (panel B), and the resulting time-by-time similarity matrix (panel C).

**Fig. 2.**
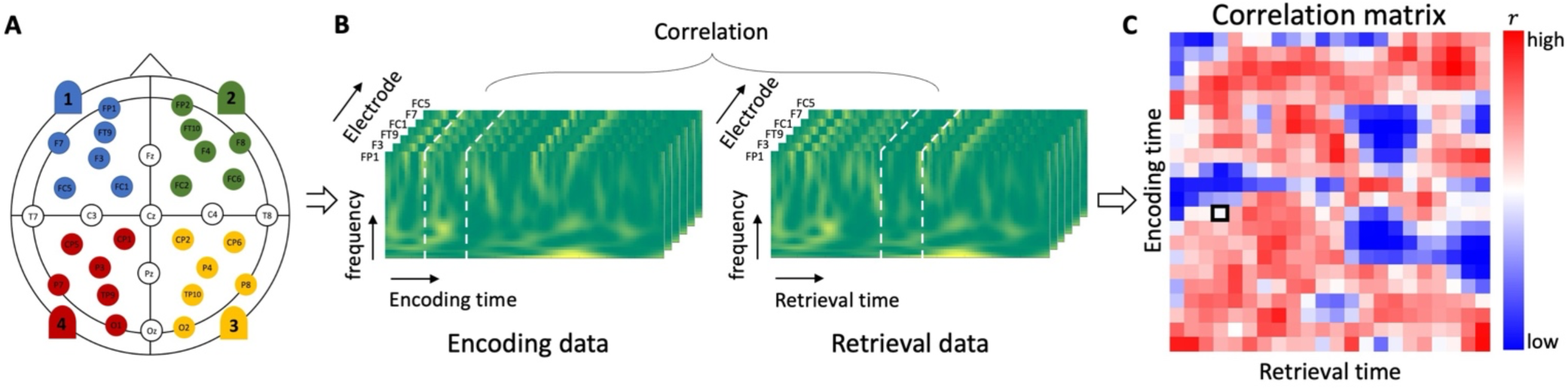
Overview of the representational similarity analysis (RSA) pipeline. (A) EEG electrodes were grouped into four non-overlapping scalp regions (left frontal, right frontal, left posterior, right posterior). (B) For each region, time–frequency power (3–40 Hz) was extracted for each 300 ms time window for each encoding and each retrieval trial. Vertical white dashed lines indicate example windows for encoding (300-600ms) and retrieval (800-1000ms) for which the log-transformed frequency values will be correlated within each electrode and then averaged across electrodes in the scalp region.(C) Encoding–retrieval similarity matrix generated by correlating encoding and retrieval feature vectors across all time-window pairs. Warm colors indicate higher similarity (r), and cool colors indicate lower similarity. The resulting time–time matrix provides a temporal map of neural reinstatement between a trial at encoding and at retrieval for all combinations of time windows within a spatial region.

For each subject, we computed within-event and between-event similarity separately for: (1) context hits—trials for which participants correctly identified a matching object–scene pairing as “same-old”; (2) context misses—trials for which participants incorrectly judged a matching object–scene pairing as “different-old”; (3) context correct rejections (CR)— trials for which participants correctly identified a mismatching object–scene pairing as “different-old”; and (4) context false alarms (FA)—trials for which participants incorrectly endorsing a mismatched object-scene pair as a previously studied match “same-old.” Separating trials according to these four recognition outcomes allowed us to quantify how strongly neural similarity patterns distinguished successful reinstatement (hits, CR) from retrieval failures (misses, FA).

For each electrode region and trial, Pearson correlations were computed between the encoding and retrieval feature vectors across all pairs of 300-ms windows, producing a two-dimensional *time × time* similarity matrix. Each matrix quantified how strongly the neural pattern during a given encoding interval was reinstated during each retrieval interval. We computed two forms of similarity: within-event similarity, reflecting correlations between a retrieval trial and its *corresponding* encoding trial (i.e., same object), and between-event similarity (B), reflecting correlations between a retrieval trial and *all other* encoding trials from the same response category (i.e. context hit, context correct rejection). Within-event similarity captured event-specific neural reinstatement, while between-event similarity served as a category-level baseline that controls for general similarities across trials of the same type.

To measure event-specific encoding-retrieval similarity, between-event similarity was subtracted from within-event similarity for each separate condition (e.g., within event context hit – between event context hit). These values were then compared across behavioral outcomes to quantify reinstatement differences associated with successful versus unsuccessful retrieval. For matching object-scene trials, memory-related reinstatement was measured by contrasting context hits and context misses:

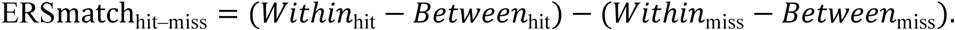

Likewise, reinstatement associated with recognition of mismatching pairs was quantified by contrasting context correct rejections and context false alarms:

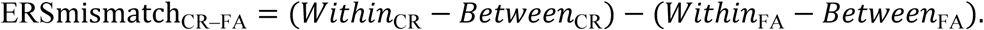

### 2.6 Multiple Regression and Cluster-Based Permutation Analysis

To evaluate how sleep characteristics relate to ERS, we fit a multiple linear regression model at each encoding × retrieval time point in the ERS matrix using MATLAB’s fitlm function. ERS values at each time–time coordinate served as the outcome variable, while age, a sleep component score, and their interaction term (age × sleep) were included simultaneously as predictors. This approach allowed us to assess both the main effects of age and sleep discontinuity on ERS as well as whether the relationship between sleep and neural reinstatement varied as a function of age. Regression coefficients were estimated independently at each time–time coordinate, producing spatiotemporal maps of the statistical effects.

To identify significant ERS temporal clusters, we first thresholded the individual encoding-retrieval time window correlations at p ≤ 0.05 (uncorrected) and then grouped them into temporally contiguous clusters. For example, if both ([200–500 ms] encoding, [500–800 ms] retrieval) and ([300–600 ms] encoding, [500–800 ms] retrieval) exceeded threshold, they were combined into a single cluster spanning ([200–600 ms] encoding, [500–800 ms] retrieval). This procedure reduces the likelihood that isolated statistical fluctuations are interpreted as meaningful effects and better captures the temporal continuity inherent in EEG data. This clustering procedure was conducted separately for each of the four electrode regions (left frontal, right frontal, left posterior, right posterior), allowing region-specific temporal patterns of sleep-related modulation of ERS to be identified without forcing spatial averaging across the scalp. Cluster significance was then evaluated using a non-parametric cluster-based permutation test. For the sleep main effect, age main effect, and Age × Sleep interaction, statistical values were permuted 10,000 times to generate a null distribution of cluster sizes. Observed clusters were considered significant if their size exceeded the cluster-size threshold derived from this permutation distribution (Maris & Oostenveld, 2007). This produced a null distribution of cluster-level test statistics, against which the observed clusters were compared. Clusters exceeding the 95th percentile of the null distribution were considered statistically significant, thus controlling for multiple comparisons across the entire time–time ERS matrix.

## 3. Results

### 3.1. Principal Components Analysis of Sleep Metrics

PCA analyses revealed two components (Table 1). The first component was characterized by high positive loadings from wake after sleep onset (WASO), number of wake bouts, and sleep fragmentation, along with a negative loading from sleep efficiency. This component was interpreted as reflecting sleep discontinuity, with higher scores indicating more fragmented and disrupted sleep. The second component was dominated by a strong positive loading from total sleep time and was interpreted as reflecting sleep time. Component scores for both principal components were extracted for each participant and used in subsequent analyses.

**Table 1.** Sleep variable loadings for the PCA of mean sleep metrics over post-encoding sleep nights.

| *Sleep Variable* | *Sleep Discontinuity (PC1)* | *Sleep Time (PC2)* |
| --- | --- | --- |
| *Wake After Sleep Onset* | **0.80** | 0.50 |
| *Number of Wake Bouts* | **0.71** | 0.59 |
| *Sleep Fragmentation* | **0.86** | - 0.001 |
| *Sleep Efficiency* | **-0.84** | 0.22 |
| *Total Sleep Time* | -0.11 | **0.96** |

*Note. The table shows the loadings for the sleep mean variables of interest. Bartlett's test confirmed adequate intercorrelations among sleep variables, $\chi^2 = 32.360$ , $df = 1$ , $p < .001$ . PCA identified two components explaining 62 % (Sleep Discontinuity) and 38% (Sleep Time) of the explained variance. Loadings are Varimax-rotated. Bold text indicates the dominant variables for each component.*
\* Co-senior authors contributed equally to this work.

### 3.2. Behavioral Results

#### 3.2.1 Sleep, context memory performance, and sleep-memory associations

Sleep discontinuity differed significantly between age groups, with older adults showing higher sleep discontinuity than younger adults (*t(68) = -3.606, p = 0.0006*). Sleep time did not differ between groups (*t(68) = -0.662, p = 0.51*) (Fig. 3A). Memory performance was quantified as the proportion of correct context responses for matching and mismatching context pairs at immediate and delayed retrieval, as well as memory retention (immediate-delayed performance). Group-level performance is illustrated in Fig. 3B. ANOVAs including factors of pair type (match, mismatch), and age (younger, older) were conducted for immediate retrieval, delayed retrieval, and memory retention separately. For immediate retrieval, there was a significant main effect of age, *F (1, 136) = 17.92, p < 0.001, ηp² = 0.12*, indicating higher performance in younger than older adults. The main effect of pair type approached significance, *F (1, 136) = 3.87, p = 0.051, ηp² = 0.03*, with higher accuracy for matching than mismatching pairs. The age × pair type interaction was not significant, *F (1, 136) = 1.76, p = 0.187, ηp² = 0.01*. For delayed retrieval, significant main effects were observed for both age *F(1, 136) = 5.01, p = 0.027, ηp² = 0.04*, and pair type, *F (1, 136) = 9.90, p = 0.002, ηp² = 0.07*. Younger adults again outperformed older adults, and matching pairs were remembered better than mismatching pairs. The interaction between age and pair type did not reach significance, *F (1, 136) = 3.56, p = 0.061, ηp² = 0.03*. For retention, there was a significant age × pair type interaction, *F(1, 136) = 14.76, p < 0.001, ηp² = 0.10*. Follow-up comparisons indicated that this interaction was driven by greater retention for mismatching pairs in older than younger adults, t (68) = −3.57, p = 0.001, d = 0.85. In contrast, there was no significant age difference in retention for matching pairs, *t (68) = 1.66, p = 0.102, d = 0.40*. Within-group analyses showed that younger adults retained matching pairs better than mismatching pairs, *t (34) = 3.41, p = 0.002, d = 1.01*, whereas older adults showed no reliable retention difference between pair types, *t (34) = −1.10, p = 0.279, d = 0.33.* Together, these findings indicate that while younger adults show superior context memory accuracy overall, age-related differences in memory retention depend on contextual congruency, with older adults showing relatively preserved retention for mismatching pairs.

**Fig. 3.**
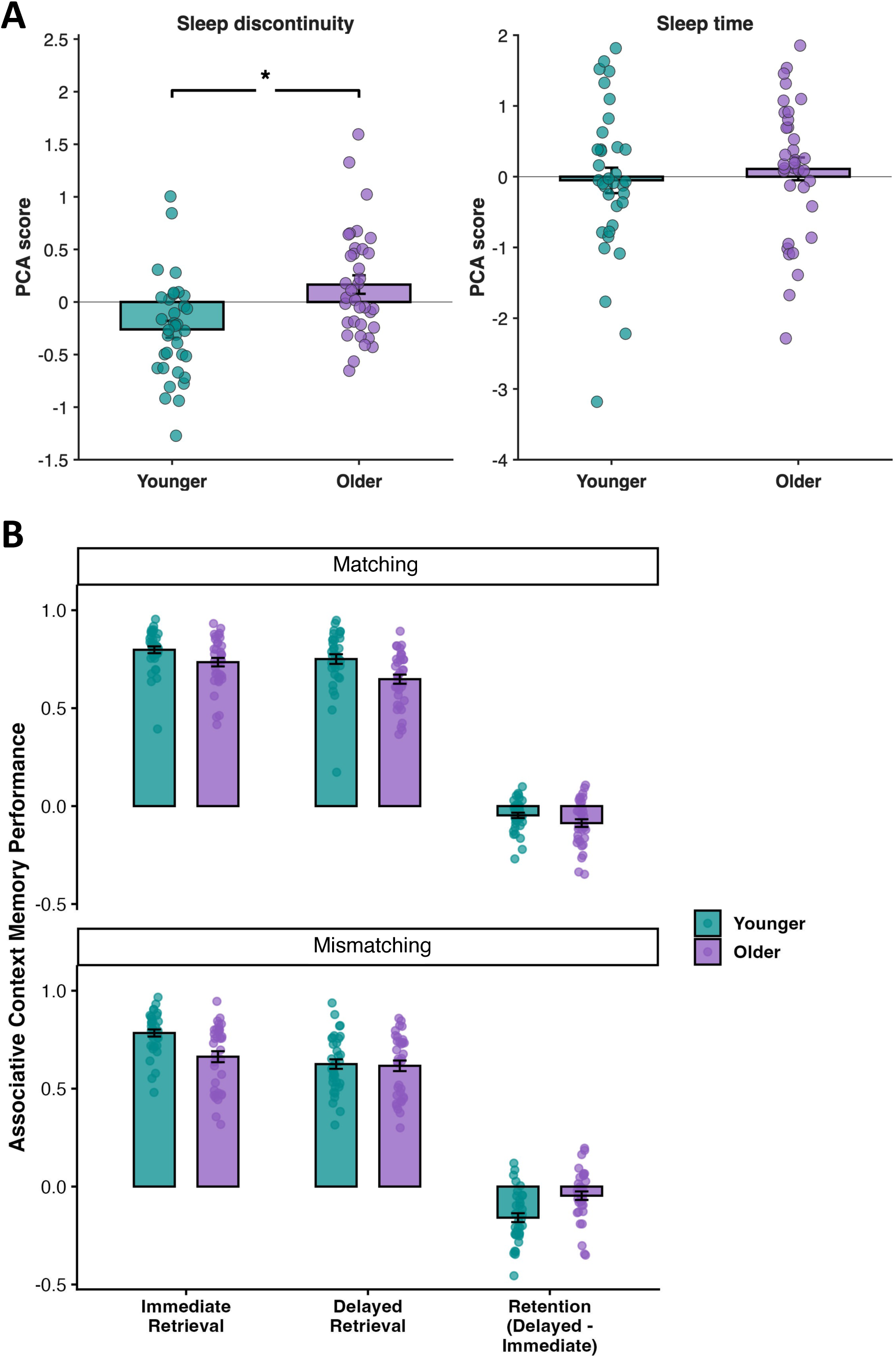
Age differences in sleep, context memory performance, and sleep-memory associations for younger and older adults. **(A)** Group distributions of sleep discontinuity and sleep time PCA scores in younger and older adults. Sleep discontinuity PCA scores differed significantly between age groups, whereas sleep time PCA scores did not. Bars represent group means ± 1 SEM, and individual participants are overlaid as points. Sleep discontinuity PCA scores differed significantly between age groups, whereas sleep time PCA scores did not. Asterisks denote statistical significance (p < 0.05). **(B)** Context memory performance is shown for younger (green) and older (purple) adults under matching (top) and mismatching (bottom) context conditions at immediate retrieval, delayed retrieval, and retention (immediate-delayed). Negative values for retention reflect the amount of forgetting relative to immediate retrieval over the delay period. Bars represent group means with ±1 SEM, dots show individual participants. Positive retention values reflect improved performance over time, whereas negative values indicate forgetting.

#### 3.2.2 Associations Between Sleep and Context Memory Performance

To examine whether individual differences in sleep during the post-encoding retention interval were associated with context memory performance, we conducted a series of hierarchical multiple regression analyses. In each model, age and retention-period sleep measures, and their interaction, were entered as predictors of delayed context memory outcomes. Separate regression models were run for each PCA-derived sleep component—sleep discontinuity and sleep time**—**to predict delayed context memory performance for matching and mismatching pairs. In all analyses, age was included as a covariate, and interaction terms were added in a second step to assess whether the relationship between sleep and memory performance differed as a function of age. Results from these models are summarized in Table 2. As can be seen in the table, only the sleep discontinuity component predicted delayed context memory accuracy for mismatching pairs across age groups, with greater discontinuity being associated with worse memory accuracy across age. Consequently, subsequent ERS analyses focus on mismatching pairs. Regression models for retention memory scores did not reveal significant associations with any of the sleep components and are reported in Supplemental Table 1. Based on these results, all subsequent analyses were limited to sleep discontinuity. Also, analyses examining pre-encoding sleep did not show the same pattern of associations with delayed context memory or ERS and are reported in the Supplemental Table 2.

**Table 2.**
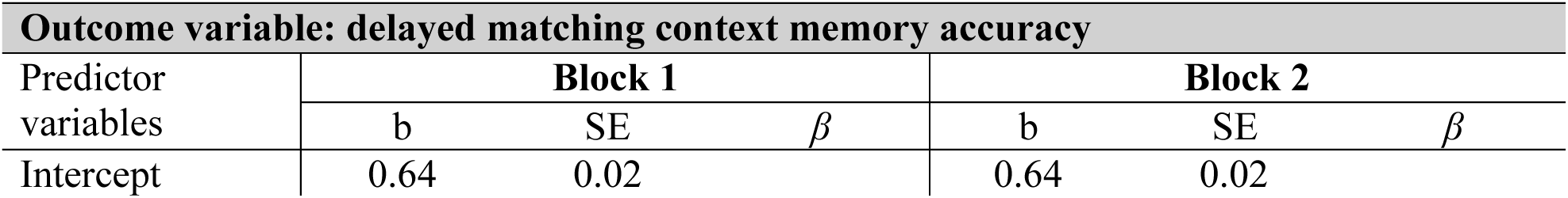
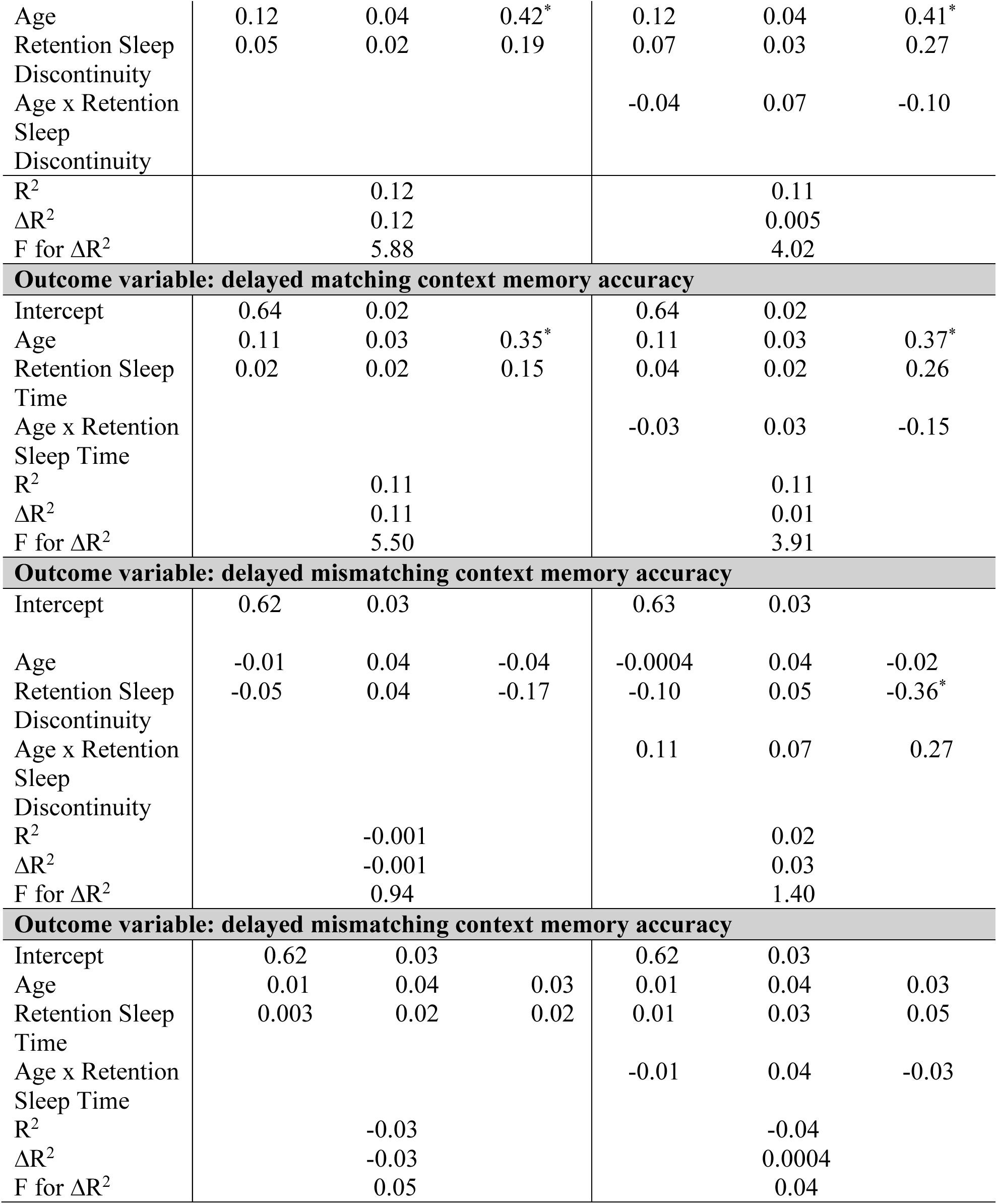
Summary of hierarchical multiple regression with age and retention sleep predicting delayed memory accuracy for matching and mismatching context pairs.

### 3.3. Associations Between Sleep Discontinuity and Encoding–Retrieval Similarity

We analyzed the relationship between sleep discontinuity and event-specific ERS for match pairs: (within trial hit – between trial hit) – (within trial miss – between trial miss); and for mismatch pairs: (within trial CR – between trial CR) – (within trial FA – between trial FA). For each trial type, we assessed the main effects of sleep discontinuity and age for ERS as well as the interaction. Given the behavioral results showed that only sleep discontinuity was associated with context memory performance for mismatch trials, we report the ERS results for mismatching context pairs. Results for matching context pairs are shown in the Supplemental Material (Supplemental Table *3, Fig. S1, Fig. S2*).

As shown in Table 3, several significant spatiotemporal clusters showed main effects of age, with older age generally associated with reduced ERS. Because significance was evaluated at the cluster level, Table 3 provides the peak-bin t-statistic and p-value within each significant cluster as descriptive summaries of the strongest effect in that cluster. For most spatiotemporal clusters, greater sleep discontinuity was associated with reduced ERS (Fig. 4). A few posterior scalp clusters showed that greater sleep discontinuity was associated with greater ERS during early encoding and late retrieval periods across age groups (Fig. 4).

**Fig. 4.**
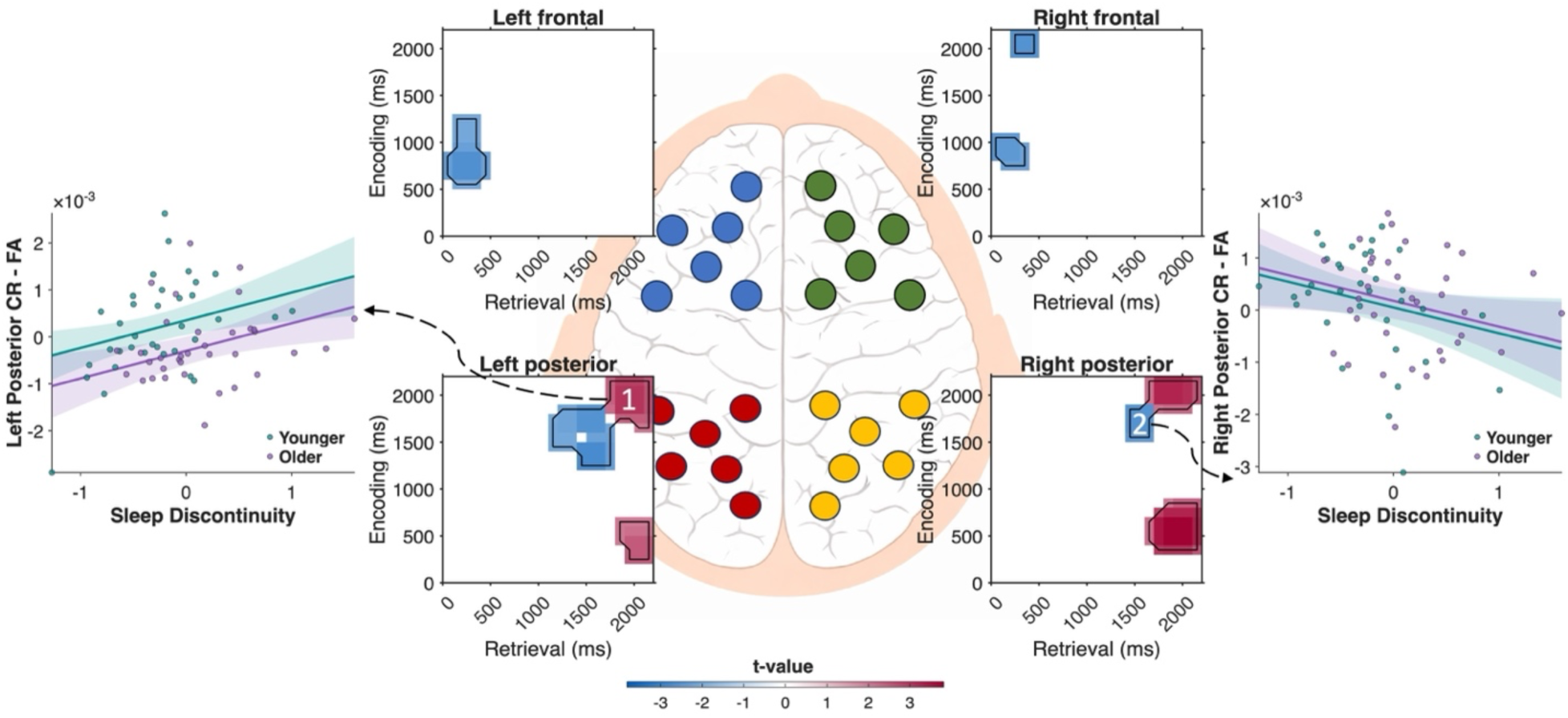
Main effect of sleep discontinuity on encoding–retrieval similarity (ERS) for mismatch context trials (CR–FA contrast), across age. Significant spatiotemporal clusters showing the association between sleep discontinuity and ERS (CR–FA) across frontal and posterior electrode regions. Insets display clusters in encoding × retrieval time space, with warmer colors indicating positive t-values and cooler colors indicating negative t-values. Two representative clusters are highlighted (Cluster 1: left posterior; Cluster 2: right posterior). Scatterplots illustrate the relationship between sleep discontinuity and ERS values for these clusters, with regression lines shown separately for younger and older adults and shaded areas representing 95% confidence intervals.

**Table 3.** Spatiotemporal clusters showing significant main effects of age, main effects of sleep discontinuity, and age × sleep discontinuity interactions on the ERS CR–FA contrast for mismatch trials. Encoding and retrieval intervals indicate the temporal extent of each cluster; t-statistics and p-values are reported for the peak bin within each cluster. Only clusters surviving multiple-comparison correction are shown.

| **Region** | **Encoding interval**<br>**(ms)** | **Retrieval interval**<br>**(ms)** | ***t*-statistic** | ***P*-value** |
| --- | --- | --- | --- | --- |
| **Main effect of Age** |  |  |  |  |
| Left frontal | 1200–1900 | 0–600 | -3.70 | <0.001 |
| Left frontal | 1200–2100 | 700–1500 | -3.23 | <0.001 |
| Left frontal | 700–1300 | 1400–2000 | 4.22 | <0.001 |
| Right frontal | 1100–1700 | 0–300 | -2.75 | <0.001 |
| Right frontal | 600–1200 | 1400–1900 | 3.25 | <0.001 |
| Left posterior | 800–1400 | 500–1000 | -2.53 | <0.001 |
| Left posterior | 0–800 | 1300–1900 | 3.22 | <0.001 |
| Left posterior | 1700–2300 | 1700–2300 | -3.81 | <0.001 |
| Right posterior | 0–300 | 200–900 | -2.76 | <0.001 |
| Right posterior | 1700–2100 | 500–900 | -2.16 | 0.020 |
| **Main effect of Sleep Discontinuity** |  |  |  |  |
| Left frontal | 500–1300 | 0–500 | -2.60 | <0.001 |
| Right frontal | 700–1100 | 0–400 | -2.39 | <0.001 |
| Right frontal | 1800–2200 | 0–500 | -3.17 | 0.001 |
| Left posterior | 1400–1900 | 1100–1700 | -2.55 | <0.001 |
| Left posterior | 1200–1700 | 1400–1800 | -2.72 | <0.001 |
| Left posterior | 1600–2200 | 1700–2200 | 3.02 | <0.001 |
| Left posterior | 200–700 | 1800–2200 | 2.47 | 0.012 |
| Right posterior | 1500–1900 | 1400–1700 | -2.53 | 0.001 |
| Right posterior | 300–900 | 1600–2200 | 3.81 | <0.001 |
| Right posterior | 1800–2200 | 1600–2200 | 3.27 | <0.001 |
| **Age × Sleep Discontinuity** |  |  |  |  |
| Left frontal* | 700–1300 | 100–700 | 3.13 | <0.001 |
| Left frontal** | 400–1200 | 700–1400 | 3.16 | <0.001 |
| Right frontal *** | 800–1300 | 0–700 | 3.17 | <0.001 |
| Right frontal* | 1800–2200 | 0–400 | 2.62 | 0.047 |
| Left posterior* | 200–600 | 0–300 | 2.22 | 0.016 |
| Right posterior* | 900–1300 | 800–1200 | 2.18 | 0.034 |
| Right posterior* | 1300–1900 | 1100–1600 | 2.48 | <0.001 |

\* Co-senior authors contributed equally to this work.
\*Significant in younger adults \*\*Significant in older adults \*\*\*Significant in both age groups

Significant age × sleep discontinuity interactions were also observed across frontal and posterior electrode regions. To interpret the age × sleep discontinuity interactions, we conducted follow-up analyses within each significant interaction cluster. ERS values were first averaged across all significant encoding–retrieval time bins within each cluster for each participant. We then tested the association between post-encoding sleep discontinuity and cluster-averaged ERS separately in younger and older adults. Follow-up results showed that most of the interactions were driven by stronger negative associations between sleep discontinuity and ERS in younger adults, across early through late encoding and retrieval windows. As shown in Fig. 5, for clusters showing significant effects in older adults over mid-to-late encoding and retrieval windows (∼400 ms encoding, ∼700ms retrieval) across frontal scalp sites, positive associations were observed, with greater sleep discontinuity corresponding to higher ERS. Together, these findings indicate that the relationship between sleep discontinuity and memory-related ERS for mismatching pairs was observed across the scalp and encoding-retrieval epochs, driven mostly by negative associations for young adults but positive associations for older adults.

**Fig. 5.**
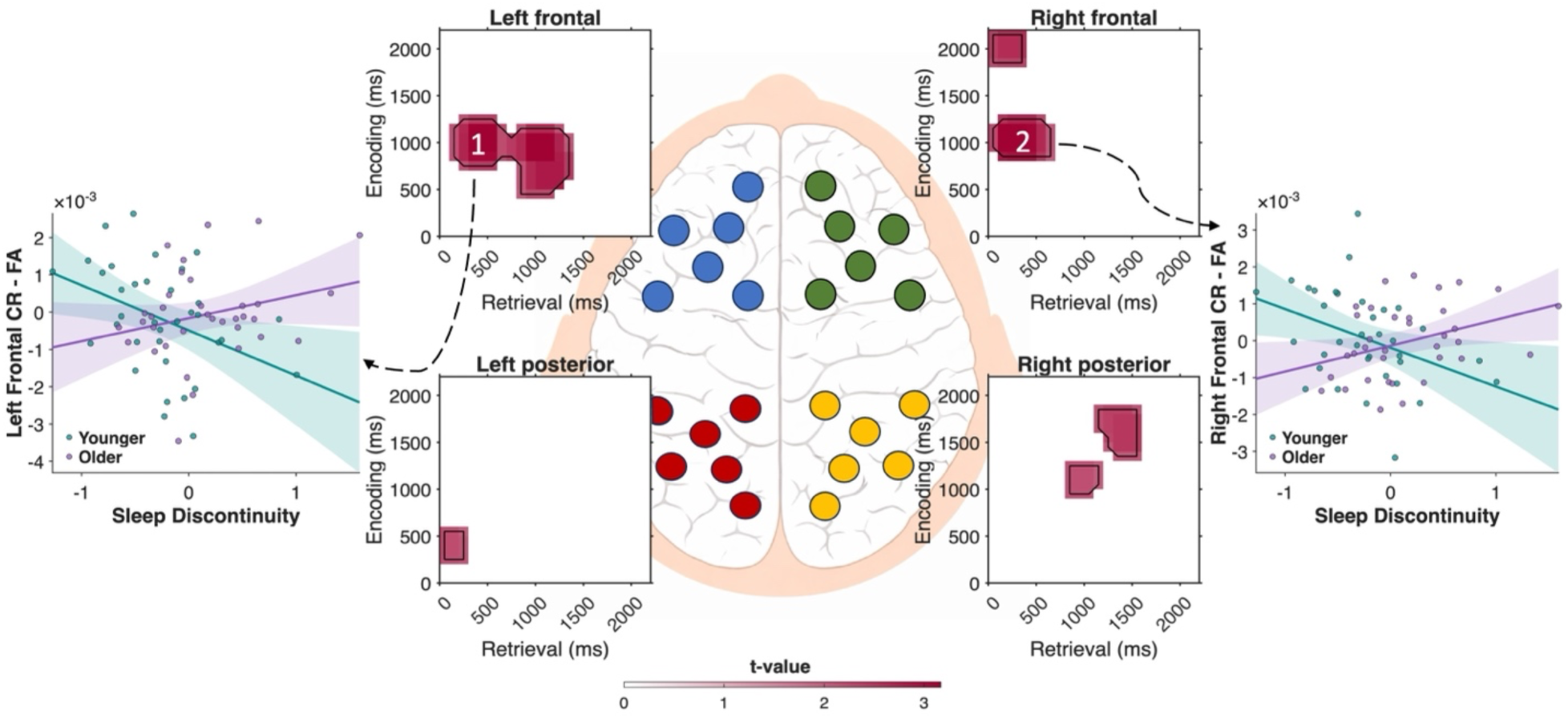
Age moderates the relationship between sleep discontinuity and encoding–retrieval similarity (ERS) for mismatch context trials. Significant age × sleep discontinuity interaction effects are shown for mismatch context memory trials. The central panel illustrates the frontal and posterior electrode groupings used in the ERS analyses. Surrounding panels depict the corresponding significant spatiotemporal clusters in encoding × retrieval time space for each region. Two representative clusters are highlighted. In Cluster 1 (left frontal), younger adults show a negative association between sleep discontinuity and the ERS CR–FA contrast, whereas older adults show a positive association. A similar pattern is observed in Cluster 2 (right frontal), where younger adults again show a negative association and older adults show a positive association. Scatterplots illustrate fitted regression lines for younger and older adults, with shaded regions representing 95% confidence intervals. Warmer colors indicate positive t-values and cooler colors indicate negative t-values in the cluster maps.

## 4. Discussion

The present study shows that post-encoding sleep quality, indexed by sleep discontinuity, is associated with delayed context memory accuracy, particularly for events dependent upon episodic reinstatement (i.e. mismatching object-context pairs), and with the neural reinstatement of encoded episodic representations during later retrieval, across age. This pattern is consistent with models proposing that sleep supports the stabilization, strengthening, and transformation of newly encoded memories, particularly for hippocampus-dependent associative information (Born & Wilhelm, 2011; Diekelmann & Born, 2010; Rasch & Born, 2013). These results extend this framework to naturalistic, multi-night sleep in younger and older adults alike, and to neural mechanisms supporting episodic reinstatement. These results and their implications are discussed below.

Older adults showed greater sleep discontinuity than younger adults, whereas sleep time did not differ significantly between age groups. Younger adults also showed better overall context memory accuracy than older adults, and matching pairs were remembered more accurately than mismatching pairs across retrieval delays and age. Correctly identifying a mismatching pair as “different old” requires more than a global sense of familiarity, because both the object and the scene are familiar at test. Instead, successful performance requires recovery of the originally studied association and use of that retrieved representation to reject the recombined probe. This interpretation closely matches recall-to-reject accounts of associative recognition and later work showing that rearranged pairs place strong demands on recollection-based discrimination (Buchler et al., 2008, 2011; Rotello & Heit, 2000). Interestingly, only sleep discontinuity—and not sleep time—predicted delayed memory accuracy for mismatching pairs. This pattern argues against a simple quantity-based account and instead supports the idea that continuity of sleep may be more critical than duration for preserving high-fidelity episodic traces. That interpretation is consistent with mechanistic accounts emphasizing coordinated offline processing during sleep and with aging models suggesting that disrupted sleep physiology may impair memory-related processing even when total sleep opportunity is not markedly reduced (Mander et al., 2017; Rasch & Born, 2013). These findings suggest that sleep quality, particularly sleep continuity, may be more relevant to ERS, and episodic memory performance, than sleep duration alone. Importantly, these associations appeared specific to the post-encoding retention interval. Exploratory analyses of pre-encoding sleep did not show the same pattern of associations with delayed context memory or ERS, suggesting that sleep after learning may be particularly important for preserving the episodic representations needed for later context discrimination. This post-encoding specificity extends prior work by isolating the retention interval more directly than studies focused on naturalistic sleep alone.

These findings replicate and extend our prior work by showing that poorer naturalistic sleep quality, including greater night-to-night variability and discontinuity, is associated with poorer episodic memory performance and reduced memory-related neural activity (Hokett et al., 2022). However, unlike Hokett et al., 2022 who reported stronger sleep–memory associations with older age, the present study showed only a trending age interaction. One possible explanation is that the current task used pictorial object–context associations, whereas prior work used verbal paired associates. Although this difference in stimulus type may have contributed to the different age-related pattern, a direct within-participant comparison of verbal and pictorial tasks would be needed to determine whether stimulus format, task demands, or differences in participant sleep and memory performance explain the discrepancy (Hokett et al., 2021, 2022; Hokett & Duarte, 2019). Moreover, differences in sample characteristics may have contributed to the weaker age moderation observed here. Prior samples included older adults with poorer sleep quality and a larger proportion of participants from racialized minority backgrounds, for whom sleep disparities are well documented (Hokett & Duarte, 2024). In contrast, older adults in the present sample may have shown relatively preserved sleep continuity, potentially reducing sensitivity to age-related differences in sleep–memory associations.

The ERS findings support the behavioral results by showing that individuals with greater post-encoding sleep discontinuity show reduced neural episodic reinstatement for correctly rejected mismatching object-context pairs during delayed retrieval. As has been shown in prior EEG and fMRI studies, ERS was also generally reduced with age, across spatiotemporal clusters, concomitant with those in memory accuracy (Chamberlain et al., 2022; Hill et al., 2020; Lee et al., 2022). Theories of sleep-dependent memory have long argued that sleep supports the strengthening and reorganization of recently encoded information, especially for associative and declarative memory (Born & Wilhelm, 2011; Diekelmann & Born, 2010; Rasch & Born, 2013). ERS frameworks, in turn, suggest that retrieval succeeds when encoded information can be re-expressed with sufficient specificity at test (Ritchey et al., 2013; Waldhauser et al., 2016; Wing et al., 2015). The present findings connect these ideas by showing that naturalistic variability in sleep continuity during the retention interval is associated with variability in reinstatement. In this sense, poorer sleep may not simply weaken memory in a global way, but may specifically degrade the quality, accessibility, or distinctiveness of the representations that must be reinstated when contextual discrimination is required.

The temporal distribution of the significant clusters further suggests that sleep and age influenced retrieval as an extended process rather than at a single isolated moment. In the current results, significant effects spanned multiple encoding and retrieval windows across frontal and posterior regions. This pattern is compatible with evidence that reinstatement-related and recollection-related neural signals can emerge rapidly after cue onset and then continue through later retrieval and monitoring operations (Hanslmayr et al., 2016; Waldhauser et al., 2016). Rather than pointing to one narrowly defined retrieval stage, the present findings suggest that fragmented post-encoding sleep may alter a broader temporal cascade through which contextual information is recovered and evaluated. This interpretation is also consistent with prior arguments that EEG studies of cognition can benefit from analyses of distributed temporal patterns, rather than relying exclusively on canonical ERP components (Bridwell et al., 2018)

One of the most interesting findings with respect to the impact of age on sleep-memory associations was that age moderated the relationship between post-encoding sleep discontinuity and ERS for mismatching pairs. In frontal scalp clusters, greater sleep discontinuity predicted lower ERS in younger adults but higher ERS in older adults. One face-value interpretation is that poorer sleep quality is associated with reduced reinstatement of specific neural representations in younger adults, but with greater ERS in older adults. This interpretation is difficult to reconcile with extant data showing similar positive, and sometimes even stronger, associations between sleep quality and episodic memory and supporting neural mechanisms across age (Hokett et al., 2021, 2022). Given the reduced sleep quality and context memory accuracy in older adults, however, it is more likely that this opposite pattern in older adults reflects compensatory recruitment or effortful engagement of post-retrieval operations that act to reinstate encoding-related representations when they are not readily recovered. The frontal scalp and late timecourse of these ERS effects are consistent with this interpretation, as ERPs associated with these strategic retrieval operations have been shown over similar scalp sites and time courses (Rugg & Curran, 2007). This interpretation is consistent with prior evidence that aging is associated with less precise reinstatement of encoded information during retrieval. In the present study, lower ERS in older adults may indicate that the neural patterns engaged during retrieval were less like those present during encoding. Such reduced encoding–retrieval overlap could contribute to poorer discrimination between matching and mismatching object–context pairs, because successful context memory requires the retrieval of specific details from the original encoding episode (Chamberlain et al., 2022; Grilli & Sheldon, 2022; Pauley et al., 2023). Overall, these results extend prior findings by showing that sleep–memory associations generalize beyond verbal paired-associate tasks to pictorial object–context memory, where successful performance requires discrimination between intact and recombined episodic details. By showing that post-encoding sleep discontinuity, but not sleep time or pre-encoding sleep, was associated with delayed memory and ERS, the present study highlights sleep continuity during the retention interval as a potentially important contributor to the preservation of high-fidelity episodic representations across adulthood.

## Conclusion

In summary, the present study demonstrates that post-encoding sleep discontinuity is associated with delayed context memory and with the neural reinstatement of encoded episodic representations during retrieval. These relationships were most evident when retrieval required rejection of familiar but recombined object–scene pairings, placing strong demands on associative discrimination and recollection-based retrieval. By showing that naturalistic sleep fragmentation during the retention interval relates both to delayed memory performance and to the temporal dynamics of ERS, the current findings extend established models of sleep-dependent consolidation into the domain of neural reinstatement. They also suggest that the neural consequences of poor sleep differ across age, highlighting sleep continuity as an important factor in understanding variability in episodic memory across the adult lifespan. Future work can build on these findings in several ways. A particularly important next step will be to combine naturalistic multi-night monitoring with polysomnography so that behavioral and ERS effects can be tied to specific physiological properties of post-encoding sleep. Such work would connect the current results more directly to mechanistic models of sleep-dependent consolidation. It will also be valuable to test whether improving sleep continuity enhances episodic neural reinstatement, and in turn, memory accuracy, especially in older adults.

## Notes

### Competing Interest Statement

The authors have declared no competing interest.

### Funding

- NIH: R21AG064309

